# SlumberNet: Deep learning classification of sleep stages using residual neural networks

**DOI:** 10.1101/2023.05.03.539235

**Authors:** Pawan K. Jha, Utham K. Valekunja, Akhilesh B. Reddy

## Abstract

Sleep research is fundamental to understanding health and well-being, as proper sleep is essential for maintaining optimal physiological function. Here we present SlumberNet, a novel deep learning model based on residual network (ResNet) architecture, designed to classify sleep states in mice using electroencephalogram (EEG) and electromyogram (EMG) signals. Our model was trained and tested on data from mice undergoing baseline sleep, sleep deprivation, and recovery sleep, enabling it to handle a wide range of sleep conditions. Employing k-fold cross-validation and data augmentation techniques, SlumberNet achieved high levels of accuracy (∼98%) in predicting sleep stages and showed robust performance even with a small and diverse training dataset. Comparison of SlumberNet’s performance to manual sleep stage classification revealed a significant reduction in analysis time (∼50x faster), without sacrificing accuracy. Our study showcases the potential of deep learning to facilitate sleep research by providing a more efficient, accurate, and scalable method for sleep stage classification. Our work with SlumberNet demonstrates the power of deep learning in sleep research, and looking forward, SlumberNet could be adapted to human EEG analysis and sleep stage classification. Thus, SlumberNet could be a valuable tool in understanding both sleep physiology and disorders in mammals.

## Introduction

Sleep research is critical for understanding health and well-being, as sleep plays a vital role in numerous physiological processes. These processes include memory consolidation, cognitive function, immune system regulation, cellular repair, and hormonal balance. Proper sleep is essential for maintaining optimal physical and mental health, and disruptions in sleep can lead to a wide range of adverse effects ^1^. Sleep disturbances, such as insomnia, sleep apnea, and circadian rhythm disorders, are increasingly prevalent in modern society, affecting millions of people worldwide ^2^. Chronic sleep disruptions have been linked to an increased risk of developing various health conditions, including obesity, diabetes, cardiovascular disease, and even certain types of cancer ^3^. By studying sleep patterns and their underlying mechanisms, we can gain valuable insights into the importance of sleep and develop novel interventions to address sleep-related disorders.

Mice serve as essential model organisms for investigating sleep patterns and their implications in humans ^4^. Sleep stages in mice can be categorized into wake, non-REM, and REM stages ^5–7^. During the wake stage, mice exhibit active brain and body functions, resulting in mixed frequency electroencephalography (EEG) signals and large electromyography (EMG) amplitudes ^8^. The non-REM stage, accounting for over 90% of sleep, is marked by cortical synchronization in the brain and a resting body, leading to lower peak EEG frequencies, higher EEG amplitudes, and smaller EMG amplitudes. The REM stage is characterized by active brain function and a motionless body, as evidenced by low EMG amplitudes ^9,10^. Classification of wake and sleep periods into these three stages based on EEG and EMG signals is a time-consuming part of sleep analysis, since many hours (or days) or data need to be analysed by visual inspection of the data.

In this paper, we present a novel deep learning model (“SlumberNet”) based on the residual network (ResNet) architecture ^11^, specifically designed for the classification of sleep states in mice using EEG and EMG signals. Our goal is to advance the understanding of sleep and its underlying mechanisms by leveraging the power of deep learning to vastly speed up sleep stage classification. We assessed both mice undergoing baseline (undisturbed sleep) and mice subjected to sleep deprivation for 12 hours. This approach allowed us to train and test the SlumberNet model in perturbed states, which, to our knowledge, has not been done before.

Deep learning has made significant strides in recent years, particularly in image classification, where convolutional neural networks (CNNs) have demonstrated exceptional performance ^12^, including in sleep stage classification using large datasets ^5^. Residual Networks (ResNet) have emerged as a prominent CNN architecture, addressing the vanishing gradient problem associated with training deeper models ^13^. Inspired by these developments, we adapted the ResNet architecture to analyze time-series data, such as EEG and EMG signals, to classify sleep states in mice. By leveraging a smaller but diverse dataset of mice, we were able to validate SlumberNet’s effectiveness and robustness against individual differences and noise.

## Results

### Data preprocessing and validation of EEG and EMG signals and spectra for training

We acquired data from a group of C57BL/6J mice in order to build a training set for the SlumberNet model. As well as being able to analyze baseline EEG and EMG signals, we wanted to increase SlumberNet’s potential utility by training it on data from animals undergoing sleep deprivation and recovery sleep. This would allow the model to handle signals from a variety of sleep paradigms that are typically incorporated into sleep analyses. Therefore, we designed a protocol encompassing all of these elements over three successive days (Fig. 1A).

**Figure 1.**
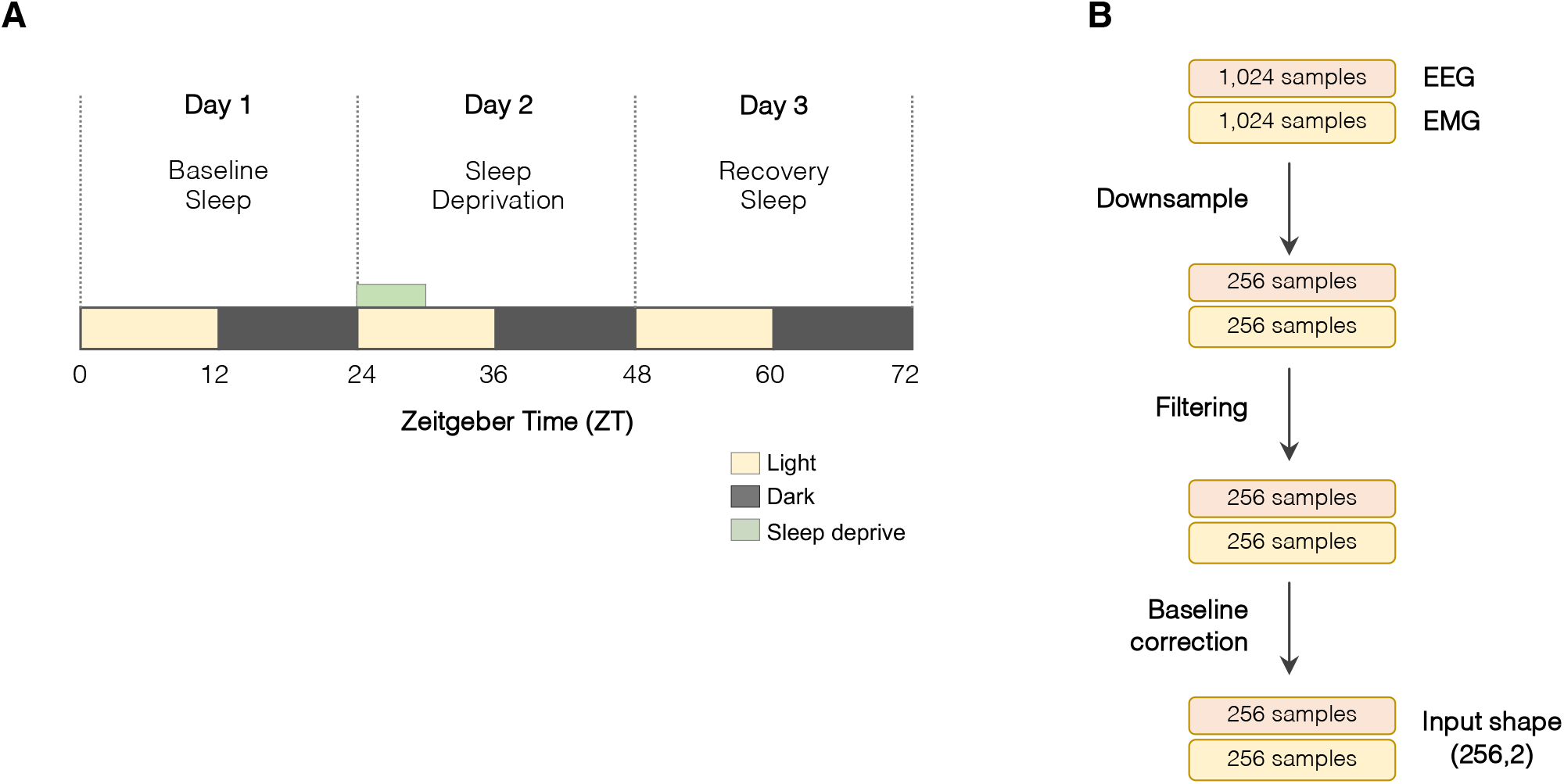
Experimental protocol, data acquisition, and preprocessing for input into SlumberNet. (A) Experimental protocol to record EEG and EMG signals for three successive days. Animals were maintained in 12h:12h light: dark cycles throughout. ZT0 indicates the onset of the first light phase; ZT12 indicates onset of the first dark phase. On the first day, animals were allowed to sleep ad libitum. On the second day, animals underwent 6 h of sleep deprivation in the first half of the light phase by gentle handling. Mice were subsequently allowed to sleep ad libitum on that day and beyond. (B) Schematic showing preprocessing steps performed on EEG and EMG data that has been sleep-staged in 4-second epochs. Data (256 Hz) were scored in 4-second epochs (1,024 samples per epoch). Paired EEM and EMG data for each epoch were downsampled to 64 Hz so that there were 256 samples per 4-second epoch. Data were filtered (EMG only) using a Butterworth filter and then baseline drift was corrected by subtracting the baseline from the raw data. The shape of the combined input array for the deep learning model was (256,2).

We acquired telemetric EEG and EMG signals (bipotential electrodes) and exported the data at a frequency of 500Hz. To classify sleep stages for the 72 h time course for each animal, we downsampled the data to 256Hz and then imported into SleepSign software for manual sleep stage classification. The data were split into 4-second epochs and each of epoch was classified as Wake (W), NREM (N), REM (R), or artefact. Artefact was attributable to movement artefact, and thus excluded from the training dataset in preprocessing (see Methods). We also excluded the entire dataset from any animal that was marred by significant electrical noise or artefact, as this was difficult to classify manually, and data from such animals would be excluded from analysis in our normal (manual) analysis workflow.

To preprocess the data for inputting into the model, we split all of the animal datasets so that we obtained matching EEG, EMG, and sleep stage data for each 4 s epoch (1,024 EEG/EMG samples at 256Hz). To decrease the total number of model parameters while retaining essential features of the data, we further downsampled the data 4-fold, such that 1,024 samples became 256 samples per 4 s epoch. The EMG signal was then put through a Butterworth filter to cut out significant electrical noise, and then both EEG and EMG signals were baseline-corrected to remove any time-dependent drift in the signals (Fig. 1B). Thus, the final electrical data that were inputted into the model consisted of an array 256 EEG and 256 EMG voltages for each 4 s epoch and a matched sleep stage for that epoch. The sleep stage data were one-hot encoded (Wake = [1,0,0]; NREM = [0,1,0]; REM [0,0,1]) and the goal of the model (its output) was to provide an array of three values (between 0 and 1) that indicates a probability given the EEG/EMG array as input.

We provide an example of 256Hz data that was used for manual sleep stage classification, and the effect of preprocessing steps on the data (Fig. 2A). The features of the data remain intact, including voltage amplitudes and frequencies, and also the spectral densities of the EEG signal (Fig. 2B). Thus, preprocessing the data for input into the model did not lead to significant loss of information used to classify the EEG and EMG signals successfully both by visual inspection and spectral metrics.

**Figure 2.**
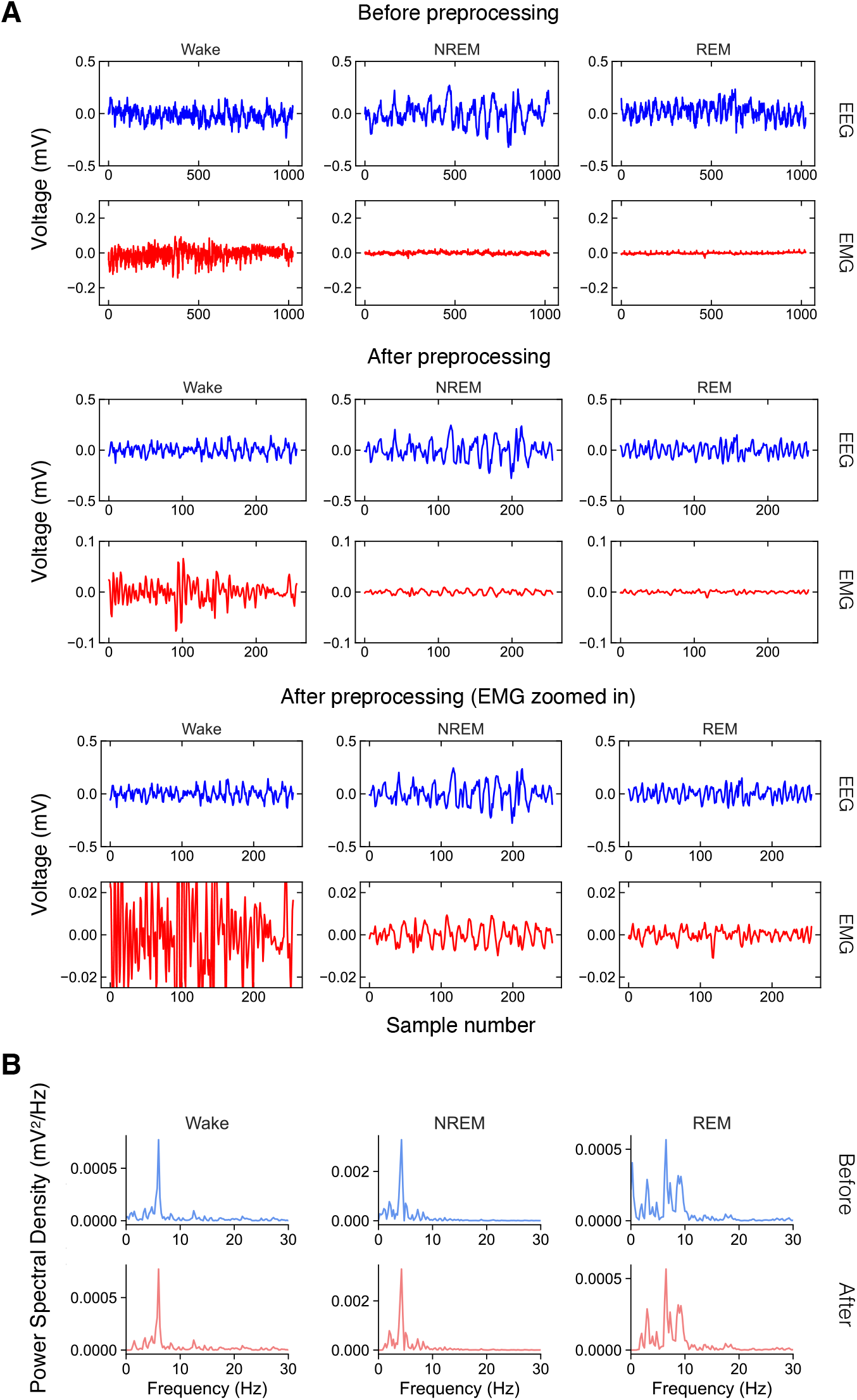
Examples of preprocessing steps for training. (A) Examples of wake, NREM, and REM EEG and EMG traces for a 4-second epoch. The traces are shown before (top) and after (middle) preprocessing has been performed. The bottom trace shows a zoomed-in y-axis for EMG traces to illustrate the amplitude differences between the three sleep stages. (B) Power spectral density (PSD) for the different sleep stages before and after preprocessing.

### SlumberNet model architecture

We built a ResNet model based on a 2-dimensional convolutional layer (Conv2D) that could take both the EEG and EMG signals as input simultaneously. The first dimension represents the time-ordered data of the voltage signal for the epoch, and the second dimension contains the signal type (EEG or EMG). This setup ensures that EEG and EMG signals are temporally matched to each other. We adopted this approach in case there are latent representations in the EMG signal that are not simply related to the absolute values (amplitude) of the EMG. Each block contains a variety of other layers including batch normalization (BatchNorm) and Dropout (Fig. 3A), although we did not need to use the dropout option as we did not observe significant overfitting in our overall model.

**Figure 3.**
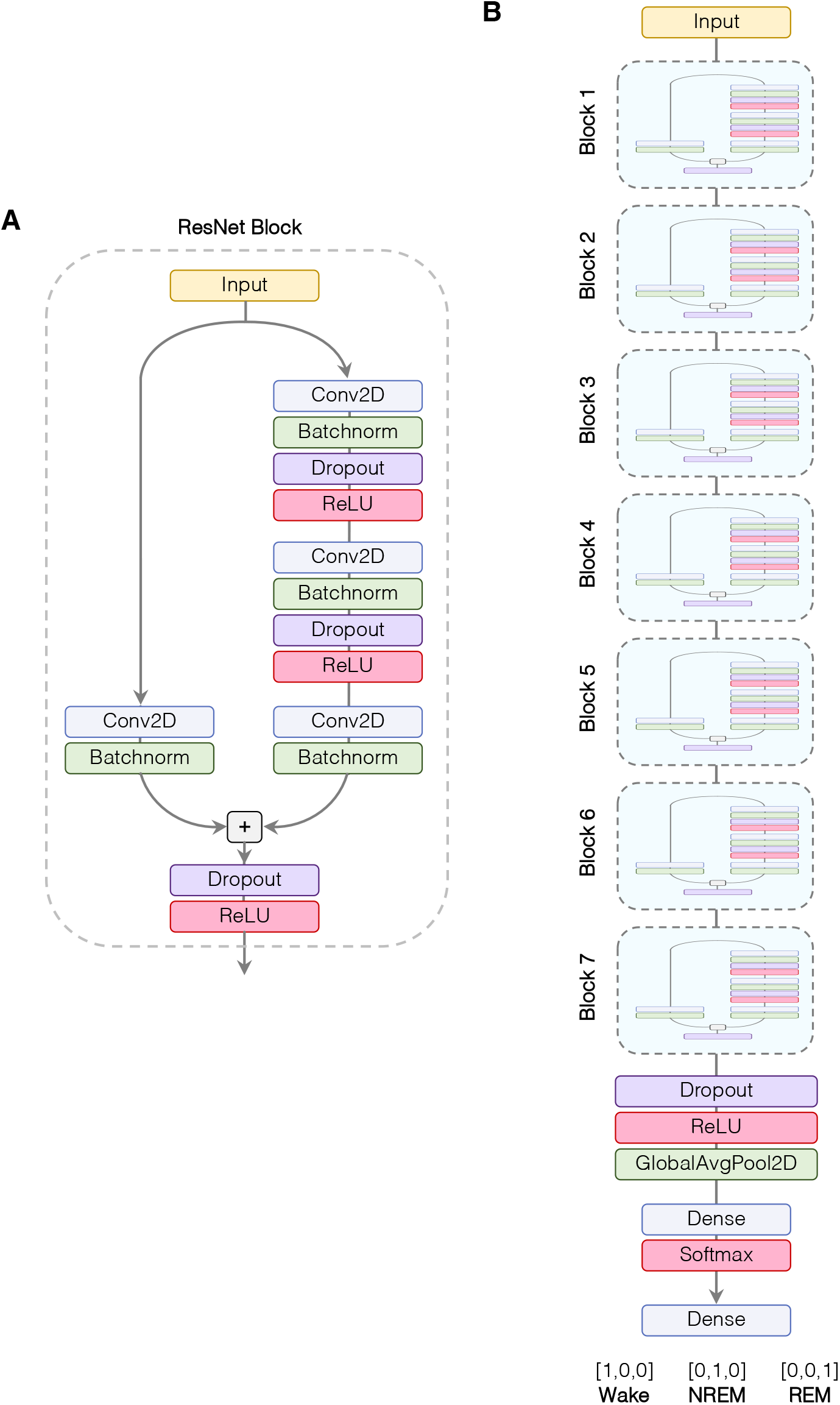
SlumberNet Model Schematic. (A) Single ResNet block architecture: Input data (2 × 256 samples: 1 × 256 EEG and 1 × 256 EMG) undergo processing by a 2D convolutional layer (Conv2D), followed by batch normalization (BatchNorm) and optional dropout. A rectified linear unit (ReLU) serves as the activation layer. This sequence is repeated before a final Conv2D layer is combined with the input via a shortcut connection in the ResNet block, ending with an activation layer. (B) Complete SlumberNet model: Input data pass through seven successive ResNet blocks with filter numbers doubling at each step. A global average pooling layer precedes a dense (fully connected) softmax-activated layer, yielding three probabilities for classification: Wake ([1,0,0]), NREM ([0,1,0]), or REM ([0,0,1]).

To produce the final model, we experimented with various numbers of concatenated ResNet blocks to balance the overall number of training parameters (which increases complexity and training time) and overfitting. We settled on seven blocks in total (Fig. 3B). This model did not overfit to the training data and thus produced validation accuracies similar training accuracies. Fewer blocks could not represent the dataset fully and accuracies failed to reach 97% (which was achieved with seven blocks). A greater number of blocks resulted in longer training times, and also overfitting to the training data without improvement in validation accuracy (which peaked at ∼97%). Such models are not useful as they will not generalize to new data that are not within the training dataset (as shown by the plateauing of validation accuracies). We did not attempt to optimize other hyperparameters (e.g. convolutional filter size, number of filters etc.).

The final layers of the model after the ResNet blocks led to global average pooling and then output of the model through a Softmax layer (normalized exponential function) to yield a final dense (fully-connected) layer of three values that represent the prediction of the sleep stage represented by the input EEG and EMG data for a 4 s epoch (Fig. 3B).

### Training and validation of SlumberNet on mouse EEG and EMG data

We designed our model to detect features purely in the EEG and EMG voltage data in a 4 s epoch, which enabled it to classify sleep stages without any information about classifications of the surrounding sleep epochs. To ensure that this setup was maintained during training, we shuffled the epochs so that the temporal order was randomized across the entire dataset. We employed a training/validation split of the training data, so that 80% of sleep samples were used for training, and the remaining 20%, which were never seen by the model for training, were used to check the accuracy of the model at each training step (training epoch).

In addition, we also used data augmentation methods to increase the generalization of the model, and to avoid overfitting. Our data augmentation involved translating the EEG and EMG data in a 4 s rolling window by a random amount. We also added random amplitude adjustments to each datapoint and added Gaussian noise. This ensured that the data fed into the model in each training epoch was different in a random way. The validation data were not augmented to ensure that the validation accuracy was consistent and comparable between training epochs.

To verify the robustness of the model we used k-fold (5-fold) cross-validation, which is a standard method used in machine learning replication ^14^. To do this, the entire dataset was split into five equal training and testing sets, such that the testing set (20% of total) each time was different to any other testing set. The remainder of sleep samples was allocated for training in each set. We ensured that there was an equal number of each type of epoch in each of the five folds (to ensure that training and testing sets were equally balanced each time). This was important since the overall proportions of Wake, NREM and REM sleep stages are different, with Wake epochs predominating. If we did not do this, the model would not generalize well because of imbalances in the training (and testing) sets between folds.

Each of the five folds was consistent, with little variability between them (Fig. 4A). In all, there was excellent agreement between training and validation accuracies (and loss), with no evidence of significant overfitting up to 50 epochs of training. Confusion matrices comparing the true and predicted sleep stages for the input data in each fold showed agreement, with the proportions of sleep stages being the same (Fig. 4B). Overall, metrics such as precision and accuracy were very similar between folds (Fig. 4C and Table 1). These data show that our model predicted Wake and NREM stages better than REM (Table 1). However, REM precision and recall were 89% (Table 1), which is comparable with other convolutional neural networks built with larger training datasets ^5^. Together, these data show that our model is capable of generalizable behavior even with a small training dataset.

**Table 1.** Accuracy, precision, and recall metrics for Wake, NREM, and REM sleep samples in each cross-validation fold. Data are percentages for each fold, and mean and standard error of the mean (SEM) data are shown for all folds.

|  | **Wake** |  |  | **NREM** |  |  | **REM** |  |  |
| --- | --- | --- | --- | --- | --- | --- | --- | --- | --- |
| **Fold** | **Accuracy** | **Precision** | **Recall** | **Accuracy** | **Precision** | **Recall** | **Accuracy** | **Precision** | **Recall** |
| 1 | 97.65% | 98.76% | 97.11% | 97.15% | 95.26% | 97.50% | 99.15% | 88.78% | 89.87% |
| 2 | 97.50% | 98.95% | 96.66% | 96.90% | 94.24% | 98.00% | 99.07% | 90.63% | 85.53% |
| 3 | 97.75% | 98.69% | 97.37% | 97.21% | 95.56% | 97.33% | 99.12% | 88.44% | 89.59% |
| 4 | 97.54% | 98.96% | 96.72% | 97.03% | 94.66% | 97.86% | 99.10% | 88.84% | 88.48% |
| 5 | 97.70% | 98.51% | 97.46% | 97.18% | 95.71% | 97.07% | 99.09% | 87.95% | 89.42% |
| **Mean** | **97.63%** | **98.78%** | **97.06%** | **97.10%** | **95.08%** | **97.55%** | **99.11%** | **88.93%** | **88.58%** |
| SEM | 0.04% | 0.07% | 0.13% | 0.05% | 0.23% | 0.14% | 0.01% | 0.37% | 0.65% |

**Figure 4.**
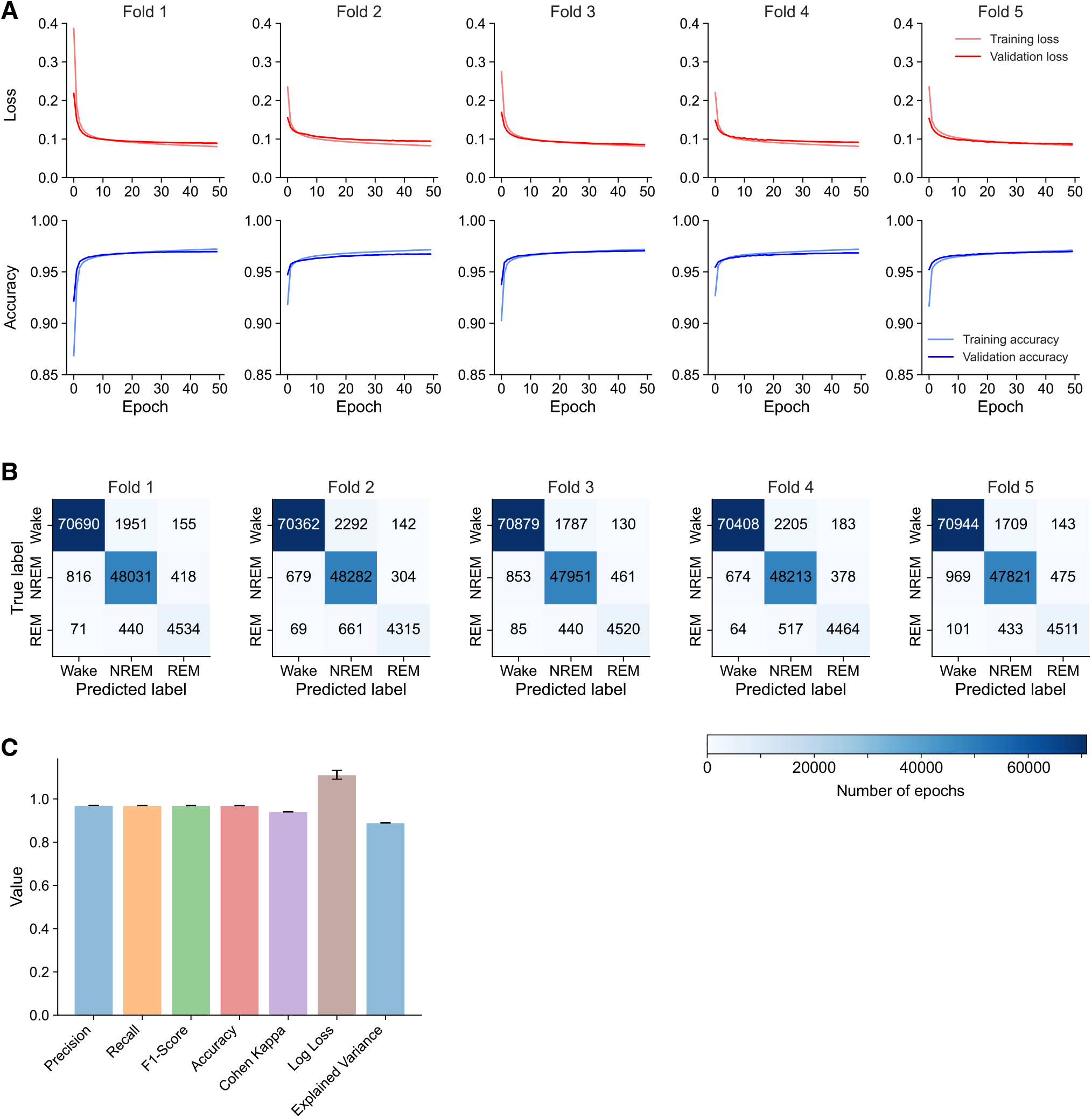
Training and validation of SlumberNet using k-fold (5-fold) cross-validation. (A) Plots showing training and validation loss and accuracy metrics for each fold of cross-validation. (B) Confusion matrices showing relationships between predicted and true labels of sleep stages. Numbers represent the absolute number of epochs assessed in the validation phase of each training fold. (C) Combined validation metrics for all folds in cross-validation (precision, recall, F1-score, accuracy, Cohen kappa, log loss and explained variance). Data are value ± SEM (n=5 training folds).

### Validation of SlumberNet on previously unseen data

To test how well SlumberNet performed on newly-acquired data, we trained the model on all of the training data. We chose the final model (after 50 training epochs) to perform further work on. We performed a new set of experiments using the same protocol as before on n=3 C57BL/6J mice (see Fig. 1A). We preprocessed data in the same way as for training, and then inputted the data into the final model and recorded the output for each sleep sample. For each 72 h time course, inference using a single graphics processing unit (GPU) took ∼1 h. In parallel, we manually scored each 4 s EEG/EMG epoch. In comparison, manual scoring for each animal took ∼ 48 h of continuous work, or ∼ 6 working days for a typical researcher.

We then compared the SlumberNet-predicted sleep stages (Wake, NREM, REM) with those obtained by using the final model’s inference (Fig. 5). There was excellent concordance between the time course data for all of the mice that we tested (Fig. 5A), with baseline sleep, sleep deprivation, and recovery sleep mapping to each other almost perfectly. Accordingly, correlation coefficients for all sleep stages were almost unity (Pearson r = 0.9984 ± 0.001; mean ± standard deviation). The only significant divergence was in the classification of Wake and NREM sleep stages, which were under- or over-predicted (respectively) by SlumberNet in Mouse 2 and Mouse 3, but not in Mouse 1 (Fig. 5A-B).

**Figure 5.**
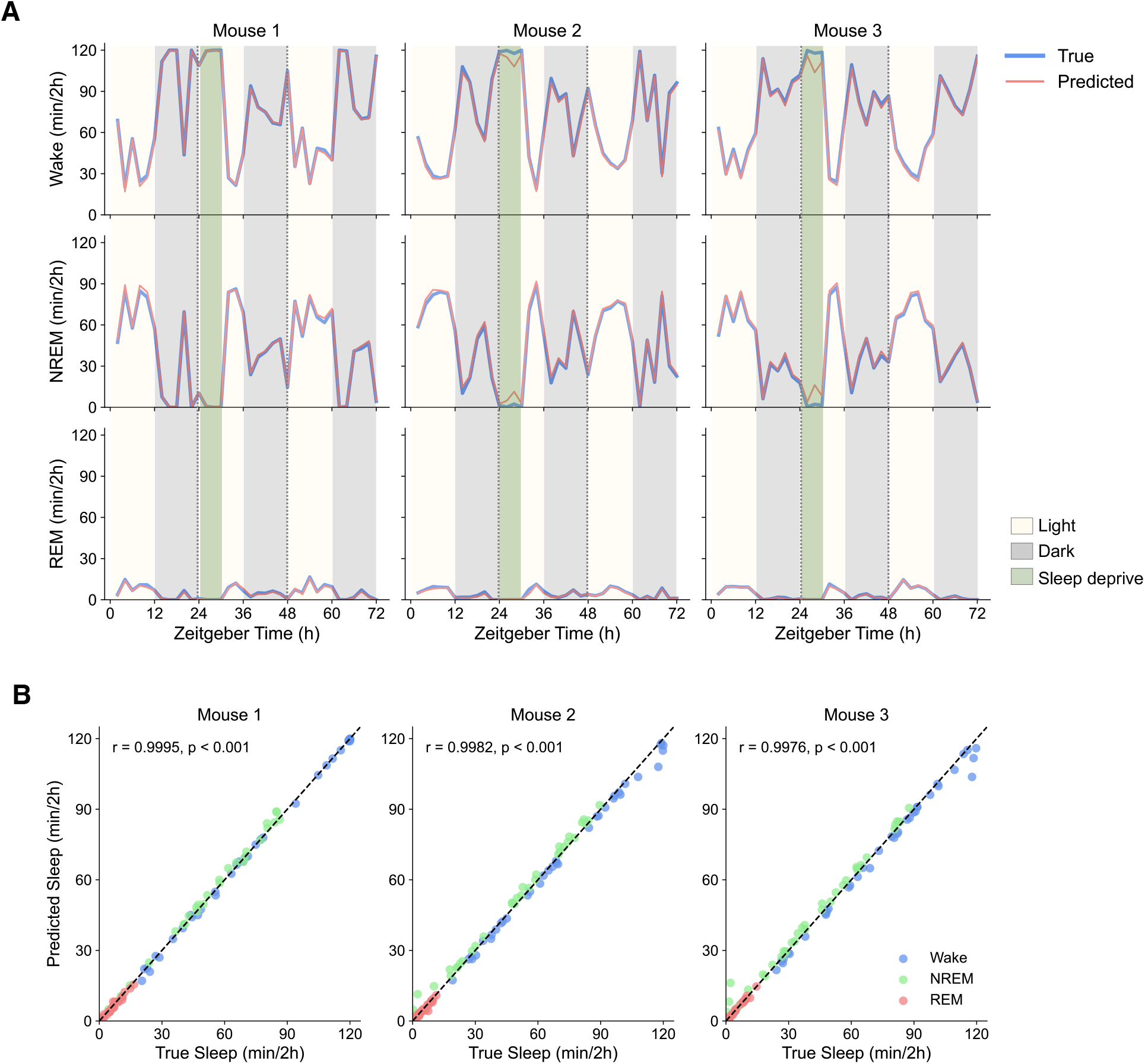
Validation of SlumberNet in experimental data not present in the training set. (A) Comparison of manually-scored sleep stage classification (True) vs model-predicted (Predicted) classification by SlumberNet. Each type of sleep stage (Wake, NREM, REM) is shown separately. The three mice were maintained in 12h:12h light:dark cycles throughout the assessments and underwent 6 h sleep deprivation on the second day (from ZT24-30; green shading). Data are plotted using 2h binning of data to calculate the number of minutes of each sleep stage per 2 h bin. (B) Correlation of True and Predicted sleep stage amount (minutes per 2 h) in each of the three mice shown in (A). Sleep stages are color-coded. Pearson correlation coefficients (r) and associated p-values are shown for each mouse.

Together, these results on independent, newly-acquired, input samples show that the final model can accurately predict sleep stages in long sleep staging time courses containing thousands of individual EEG/EMG epochs. Moreover, given the time saved by using the model (∼50x faster, or ∼150x faster accounting for the total number of days it would take manually), SlumberNet should make sleep staging far more rapid than manual approaches, without sacrificing accuracy.

## Discussion

Our study presents SlumberNet, a deep learning model based on the ResNet architecture, specifically designed for the classification of sleep states in mice using EEG and EMG signals. Our model demonstrated remarkable accuracy in classifying sleep stages, even when trained on a smaller dataset compared to previous models. This highlights the potential of SlumberNet for advancing our understanding of sleep patterns and their underlying mechanisms, as well as for facilitating future research into sleep disorders and sleep-enhancing interventions. Importantly, SlumberNet was capable of generalizing to previously unseen data, showing strong concordance between model-predicted sleep stages and manual sleep stage classification, while significantly reducing the time required for sleep analysis.

One potential future application of SlumberNet is its adaptation for human EEG analysis in sleep studies. Given the similarities between sleep patterns in mice and humans ^15^, it is conceivable that the same deep learning approach used in SlumberNet could be applied to human sleep data. This could enable more accurate and efficient analysis of human sleep patterns, aiding in the development of personalized sleep interventions and improving our understanding of sleep disorders. Furthermore, the successful adaptation of SlumberNet to human sleep data could pave the way for the development of real-time sleep monitoring and intervention systems, which would be invaluable in both clinical and research settings.

However, our study has some limitations that should be acknowledged. First, the SlumberNet model may struggle to classify sleep stages in the presence of very noisy data, such as those marred by significant electrical noise or artefacts. This limitation could be addressed by developing preprocessing techniques that effectively remove noise and artifacts from the data, thus improving the model’s performance. Additionally, it should be noted that our model was trained and tested on a relatively small dataset, which may limit its generalizability to larger and more diverse datasets. Future studies should seek to train and validate SlumberNet on larger datasets, encompassing a wider range of species and experimental conditions, in order to further enhance its robustness and applicability.

Compared to previous methods, SlumberNet offers several advantages. Firstly, the adaptation of the ResNet architecture for time-series data allows for more effective analysis of EEG and EMG signals, as well as better handling of the vanishing gradient problem associated with training deeper models ^11^. This results in improved classification performance and reduced training times. Secondly, the use of data augmentation techniques and cross-validation ensures the robustness of our model, minimizing overfitting and allowing for better generalization to new data. Lastly, the application of our model to sleep-deprived mice demonstrates its versatility and applicability to a wide range of sleep paradigms. In conclusion, SlumberNet represents a step forward in automated sleep stage classification, with the potential to accelerate our understanding of sleep patterns, disorders, and interventions.

## Methods

### Animals

Animal studies adhered to approved guidelines from the Institutional Animal Care and Use Committee (IACUC) at the Perelman School of Medicine at the University of Pennsylvania, and ARRIVE guidelines. We used wild type, male C57BL/6J mice from Jackson Laboratories and allowed them to acclimate for a minimum of two weeks before experiments. For sleep deprivation experiments, mice were housed individually in automated sleep fragmentation chambers (Model #80391, Campden/Lafayette Instrument Lafayette, IN, USA). At all times, the mice were given *ad libitum* access to food and water under standard housing conditions, under a 12-hour light: 12-hour dark cycle.

### Sleep Recording and Data Acquisition

We implanted telemetry transmitters (HD-X02, Data Sciences International, St. Paul, MN, USA) connected to electrodes for continuous EEG/EMG recording in mice aged between 9-11 weeks. The mice were anesthetized with isoflurane (induction 3-4%, maintenance 2-2.5%), and two stainless steel EEG electrodes (length of screw shaft: 2.4 mm; head diameter: 2.16 mm; shaft diameter: 1.19 mm; Plastics One, Roanoke, VA, USA) were implanted epidurally over the right frontal and parietal cortices. The electrodes were connected to the telemetry transmitter with medical-grade stainless steel wires and secured with dental cement (Kemdent, Purton, Swindon, UK). Two EMG stainless-steel leads were inserted into the neck muscles ∼5 mm apart and sutured in place. The telemetry transmitter was positioned in a subcutaneous pocket on the left dorsal flank. We administered analgesia during the surgery (subcutaneous injection of buprenorphine (Vetergesic) at 0.1 mg/kg and meloxicam (Metacam) at 10 mg/kg) and allowed animals to recover for at least 10 days before starting experimental protocols. We then recorded EEG/EMG signals continuously for 6-7 days using Data Sciences International hardware and Dataquest ART software (Data Sciences International, St. Paul, MN, USA) with a 500 Hz export rate for downstream analysis.

### Sleep stage scoring

Data were downsampled to 256Hz from 500Hz and vigilance states were determined using SleepSign for Animal ver. 3 (Kissei Comtec, Nagano, Japan) as detailed previously ^16^. Briefly, we performed manual sleep stage scoring in 4-second epochs by waveform and FFT spectrum recognition. Low amplitude EEG and high amplitude EMG signals were considered as Wake. Slow waves and high amplitudes of EEG coupled with low amplitude EMG signals were considered NREM. Low amplitude EEG dominated by theta frequencies (5–9 Hz), and loss of EMG muscle tone was defined as REM. Defined sleep-wake stages were cross-examined and corrected if necessary.

### Sleep Deprivation

Sleep deprivations were performed using a device that applied tactile stimulus with a horizontal bar sweeping just above the cage floor (bedding), as described previously ^16^. Once the sweeper was on, animals needed to step over it to continue their normal activities. Sleep deprivation began at the start of the light cycle (Zeitgeber Time 0 (ZT0)) and lasted for 12 hours, with continuous sweeping mode (approximately 7.5 seconds cycle time). We made additional attempts to maintain wakefulness during the second half of the sleep deprivation period by occasionally tapping on the cage or gently touching the animals with a brush. To evaluate the effects of 12-hour sleep deprivation on sleep/wake behaviors, we recorded baseline EEG/EMG data for three days after mice had acclimated for a week. Mice were then recorded for an additional 3-4 days during the sleep deprivation and recovery phases. After sleep deprivation, animals were allowed to recover for 24 hours.

### Data Preparation

Initially, we gathered EEG and EMG information from several files, each containing paired raw voltage data and manually identified sleep stages. The raw voltage data included EEG and EMG signals sampled at a rate of 256 Hz. We arranged these files in a Pandas dataframe, ensuring they were in the appropriate format with both voltage columns represented by floating point numbers. Files not conforming to this format were excluded from further analysis. Upon loading the voltage data, we extracted sleep stage labels from the matched sleep scoring files, designating sleep stages as wake (W), non-REM sleep (N), or REM sleep (R). Any other labels in the epoch files were considered artifacts and labeled as “A”. We then iterated through the epochs, extracting corresponding EEG and EMG voltage data and adding it to the overall data array. Concurrently, we created a matching NumPy array for the epoch labels, employing one-hot encoding for the sleep stages (W: [1,0,0], N: [0,1,0], R: [0,0,1], A: [1,1,1]).

Next, we downsampled the EEG and EMG data and conducted baseline correction for each epoch. The original EEG and EMG data consisted of 1024 samples per epoch, which we reduced by a factor of 4 to achieve 256 samples per epoch. This downsampling was vital for decreasing the computational load while preserving sufficient data for precise sleep stage classification. We utilized the resample_poly method from the SciPy library, which combines polyphase filtering and resampling, to perform the downsampling. This method is preferred over alternatives such as decimate or resample, as it provides superior anti-aliasing properties and enhanced signal preservation. We then applied a high-pass filter to the downsampled EMG signals to eliminate slow drifts and further minimize noise. The filter’s cutoff frequency was set at 0.5 Hz, as it effectively removed slow drifts without significantly altering essential signal features. We used the Butterworth filter from the SciPy library for this task, as it provides a smooth, distortion-free response in the passband. Each 4 second epoch data for EEG and EMG were then baseline corrected. Lastly, we discarded epochs labeled as artifacts (A) from both the voltage and epoch arrays. The resulting preprocessed EEG and EMG data, along with the corresponding epoch labels, were saved for later use in training the model.

### Model Training and Evaluation

We used Keras and Tensorflow2 to construct and train our model. To guarantee reproducibility, we set a random seed and leveraged all available GPUs on our server via the MirroredStrategy approach for distributed training on multiple GPUs. We employed mixed precision computing (float16 and float32) to accelerate training on Nvidia GPUs with a compute capability of 6.0 or higher, as this method uses lower-precision data types for quicker computations without sacrificing accuracy.

We built our 2D ResNet model with seven ResNet blocks, eight feature maps (kernels), and kernel dimensions of (2, 1). Although dropout was an option the ResNet blocks, we set the dropout rate to zero, as we did not observe overfitting. Dropout was also set to zero elsewhere. We employed data augmentation to increase the diversity of the training data, enhancing the model’s generalization capabilities. We implemented an AugmentDataGenerator class with a custom augmentation function for TensorFlow-based data augmentation during training. The function applied amplitude scaling to the input EEG and EMG data independently, and both signals were translating by a random amount so that they were temporally yoked together. Finally, a low-level of Gaussian noise was added to the EEG and EMG independently. These transformations helped the model to become more robust against variations in signal strength and to learn invariant features across different time shifts. As a result, the risk of overfitting was minimized.

For training, we subjected our dataset to 5-fold cross-validation, a technique for evaluating a model’s performance by dividing the dataset into five subsets or “folds.” The model was trained over 50 epochs with a learning rate of 1e-06 and a per-GPU batch size of 128. We used the Adam optimizer, an adaptive learning rate optimization algorithm, and wrapped it with a Loss Scale Optimizer to avoid multi-GPU crashes caused by underflow in low-precision computations.

We then established a Stratified Shuffle Split for k-fold cross-validation, a technique that maintains the original class distribution within each fold (original proportions of Wake, NREM, and REM classes within each fold and within the training and validation splits), ensuring that each subset is representative of the overall class distribution. This method provides a more robust evaluation of the model’s performance. Using the k-fold cross-validation technique, we trained our model and saved the best model for each fold with a ModelCheckpoint callback, which saves the best version of the model based on a specified metric. A ReduceLROnPlateau callback was also applied to decrease the learning rate if training loss did not improve for three consecutive epochs, enabling more fine-grained optimization during training.

### Performance metrics

We loaded the best model for each k-fold cross-validation fold, predicted test set class labels, and calculated various evaluation metrics, such as precision, recall, F1-score, accuracy, confusion matrix, kappa, loss, and explained variance, as outlined below.

Precision is a measure of the accuracy of a model’s positive predictions, calculated as the proportion of true positives out of the total predicted positives. It is a useful metric when the cost of false positives is high.

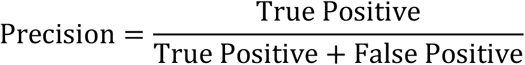

Recall is a measure of a model’s ability to identify all relevant instances, calculated as the proportion of true positives out of the total actual positives. It is useful when the cost of false negatives is high.

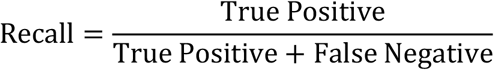

The F1-score is a combined metric that balances both precision and recall, giving equal weight to both metrics. It is calculated as the harmonic mean of precision and recall, and provides a good balance between the two metrics when optimizing for both is important.

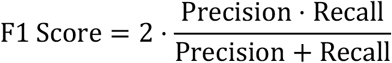

Accuracy is a measure of the overall correctness of a model’s predictions, calculated as the proportion of correct predictions out of the total number of predictions. It is a simple and intuitive metric, but can be misleading when the classes are imbalanced.

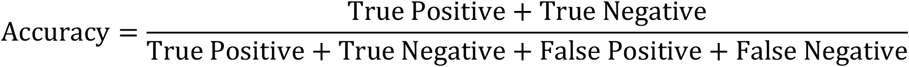

Kappa is a metric that measures the agreement between a model’s predictions and the true values, taking into account the expected probability of agreement due to chance. It is a useful metric when the classes are imbalanced or when evaluating inter-rater agreement. A Kappa > 0.8 suggests that the scoring results are in nearly perfect agreement ^5^.

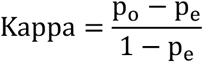

where *p*_0_ is the observed agreement and *p*_*e*_ is the chance agreement.

Loss is a metric that quantifies the difference between a model’s predicted output and the actual output for a given input. It is typically used during the training phase to optimize the model’s parameters by minimizing the value of the loss function.

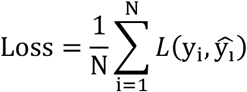

where *N* is the number of samples, _*Yi*_ is the true value of the *i*th sample, and 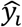 is the predicted value of the

*i*th sample, and *L* represents the loss function used in the deep learning model. We used Categorical Cross-Entropy for our loss function, which is defined by:

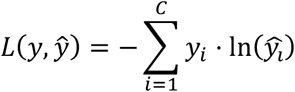

where _*Y*_ is the true label for a sample, 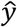 is the predicted probability distribution over the possible classes, and *C* is the number of classes.

The categorical cross-entropy loss function is commonly used in multi-class classification problems like ours. The goal of the model is to predict the probability distribution over the possible classes, and the loss function measures the difference between the predicted distribution and the true distribution. The negative sign in the equation ensures that the loss is always a positive value. The smaller the value of the loss, the better the model is performing on the training data.

Explained variance is a metric that measures the proportion of the variation in the true values that is explained by the variance in the predicted values. It is a useful metric when evaluating the performance of a regression model, and provides insight into how well the model fits the data.

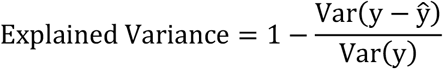

where _*Y*_ is the true value and 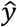 is the predicted value.

### Computer hardware

We used a custom-built server for preprocessing and model training and testing:

- Intel i9-7980XE CPU @ 2.60GHz (16-core)
- 128GB System Memory (DDR4 3000 MHz)
- 3x Nvidia Titan V (GV100) GPUs 12 GB VRAM each
- 2048GB Samsung SSD 850

The server was running on Linux (Ubuntu 22.04 LTS), Nvidia Driver Version: 525.78.01, CUDA Version: 12.0, Python 3.10.10, Tensorflow 2.11.0

## Acknowledgements

A.B.R. acknowledges funding from the Perelman School of Medicine, University of Pennsylvania, the Institute for Translational Medicine and Therapeutics (ITMAT) at the University of Pennsylvania. This work was supported also by NIH DP1DK126167 and R01GM139211 (A.B.R.).

## Author contributions

P.K.J. performed animal experiments and manual sleep stage scoring. U.K.V. analyzed experimental results and designed and implemented model code. A.B.R. designed the project and the ResNet model, analyzed results, secured funding, and wrote the paper with contributions from the other authors.

## Data availability

The datasets analyzed during the current study are available from the corresponding author on reasonable request.

## Code availability

The code for preprocessing and model definition training and testing will be made available on Zenodo.

